# Coherence of *Microcystis* species revealed through population genomics

**DOI:** 10.1101/541755

**Authors:** Olga M. Pérez-Carrascal, Yves Terrat, Alessandra Giani, Nathalie Fortin, Charles W. Greer, Nicolas Tromas, B. Jesse Shapiro

## Abstract

*Microcystis* is a genus of freshwater cyanobacteria which causes harmful blooms in ecosystems worldwide. Some *Microcystis* strains produce harmful toxins such as microcystin, impacting drinking water quality. *Microcystis* colony morphology, rather than genetic similarity, is often used to classify *Microcystis* into morphospecies. However, colony morphology is a plastic trait which can change depending on environmental and laboratory culture conditions, and is thus an inadequate criterion for species delineation. Furthermore, *Microcystis* populations are thought to disperse globally and constitute a homogeneous gene pool. However, this assertion is based on relatively incomplete characterization of *Microcystis* genomic diversity. To better understand these issues, we performed a population genomic analysis of 33 newly sequenced genomes (of which 19 were resequenced to check for mutation in culture) mainly from Canada and Brazil. We identified eight *Microcystis* clusters of genomic similarity, only four of which correspond to named morphospecies and monophyletic groups. Notably, *M. aeruginosa* is paraphyletic, distributed across four genomic clusters, suggesting it is not a coherent species. Most monophyletic groups are specific to a unique geographic location, suggesting biogeographic structure over relatively short evolutionary time scales. Higher homologous recombination rates within than between clusters further suggest that monophyletic groups might adhere to a Biological Species-like concept, in which barriers to gene flow maintain species distinctness. However, certain genes – including some involved in microcystin and micropeptin biosynthesis – are recombined between monophyletic groups in the same geographic location, suggesting local adaptation. Together, our results show the importance of using genomic criteria for *Microcystis* species delimitation and suggest the existence of locally adapted lineages and genes.

**Importance:** The genus *Microcystis* is responsible for harmful and often toxic cyanobacterial blooms across the world, yet it is unclear how and if the genus should be divided into ecologically and genomically distinct species. To resolve the controversy and uncertainty surrounding *Microcystis* species, we performed a population genomic analysis of *Microcystis* genome from public databases, along with new isolates from Canada and Brazil. We inferred that significant genetic substructure exists within *Microcystis*, with several species being maintained by barriers to gene flow. Thus, *Microcystis* appears to be among a growing number of bacteria that adhere to a Biological Species-like Concept (BSC). Barriers to gene flow are permeable, however, and we find evidence for relatively frequent cross-species horizontal gene transfer (HGT) of genes that may be involved in local adaptation. Distinct clades of *Microcystis* (putative species) tend to have distinct profiles of toxin biosynthesis genes, and yet toxin genes are also subject to cross-species HGT and local adaptation. Our results thus pave the way for more informed classification, monitoring and understanding of harmful *Microcystis* blooms.

## Introduction

Several categories of species concepts can be used as the basis to delimit and organize biological diversity, and two concepts in particular have been recently applied to bacteria and archaea. The Ecological Species Concept (ESC) proposes that speciation is driven by divergent natural selection between distinct ecological niches, while the Biological Species Concept (BSC) emphasizes gene flow (e.g. homologous recombination) as a cohesive force within species (1). The Stable Ecotype Model (SEM) is a version of the ESC tailored to bacteria, under the general assumption that adaptive mutations spread more rapidly by clonal expansion than by recombination (2). However, certain bacteria and archaea have relatively high rates of recombination, such that a BSC-like concept could apply - but strictly the BSC will never apply to bacteria that reproduce clonally and occasionally exchange genes across species boundaries. Several archaea and bacteria appear to fit a BSC-like model, showing higher recombination within species than between species (1, 3–5). Barriers to recombination could be maintained by natural selection or genetic incompatibilities, or due to physical separation (*i.e*. allopatry). Allopatric speciation is thought to be rare for globally dispersed bacteria, but does appear to occur among geographically separated hotspring archaea (6). Therefore, a plurality or spectrum of species concepts is probably necessary to fit the diverse lifestyles and recombination frequencies observed across microbes (1).

When two different species inhabit a common niche or geographic location, they may exchange genes beneficial to local adaptation. For example, *Vibrio cholerae* ‘core’ housekeeping genes are freely recombined among *V. cholerae* from both Bangladesh and the American coast, to the exclusion of the sister species *V. metecus* (7). However, in the integron (part of the genome subject to particularly frequent recombination), *V. cholerae* undergoes more genetic exchange with *V. metecus* from the same geographic location (USA) than with *V. cholerae* from a different location (Bangladesh). This suggests that species cohesion is maintained across most of the core genome, while certain ‘accessory’ genes are exchanged across species boundaries to promote local adaptation. Identifying such locally adapted genes that cross species boundaries can provide insight into the genetic basis of adaptation to different environments (8, 9).

Here we consider the common bloom-forming cyanobacterium *Microcystis* as a model of speciation and local adaptation. *Microcystis* is a genus containing a great deal of genetic diversity (10) and capable of frequent recombination (11–13). Previous genetic and genomic studies have suggested that *Microcystis* is globally distributed, with little geographic structure (10, 11, 14, 15). Thus, it is plausible that *Microcystis* represents a single, globally distributed and homogeneous gene pool, adhering to a BSC-like model (1, 5, 11, 16). However, multiple attempts have been made to classify *Microcystis* into several species, based on various morphological and genetic criteria (17–19).

*Microcystis* is able to form colonies or cell aggregates covered by exopolysaccharide or mucilage (20). Several *Microcystis* strains are known to synthesize intracellular toxins, which are thought to be released to the environment primarily when cells die and lyse (21). Together, toxins and cell decomposition followed by oxygen depletion threaten the health of humans and animals. *Microcystis* colony morphology, cell size, and the structure of mucilage have been used for decades as taxonomic criteria to classify *Microcystis* in morphospecies or morpho-types (International Code of Botanical Nomenclature) (17, 22). However, morphospecies classifications are often inconsistent with genetic, genomic, and phylogenetic analyses (17, 23). These inconsistencies may occur because *Microcystis* colonies can change morphology or become unicellular without any genotypic changes (17, 20, 24–26).

At least 51 morphospecies have been described within the *Microcystis* genus (http://www.algaebase.org/browse/taxonomy/?id=7066) (*e.g. M. aeruginosa, M. panniformis, M. viridis, M. wesenbergii, M. flos-aquae, M. novacekii*, and *M. ichthyoblabe*) (17, 27) One of the most studied and frequently reported is *M. aeruginosa*. Under laboratory conditions, it has been shown that colonies of *M. wesenbergii* morphospecies could become morphologically similar to colonies of *M. aeruginosa*, after just a few hours of culture (26). If a *Microcystis* strain undergoes colony morphology changes, it can then become over-classified into several morphospecies. Thus, the number of morphospecies may not reflect the number of *Microcystis* species based on other bacterial systematic approaches (17).

Because of these inconsistencies, several authors have attempted to reclassify *Microcystis* morphospecies using additional systematic approaches, like 16S rRNA gene sequence identity, DNA-DNA hybridization, phylogenetic analysis of conserved genes, and Average Nucleotide Identity (ANI) (10, 12, 14, 17, 19, 28, 29). For example, five *Microcystis* morphospecies (*M. aeruginosa, M. ichthyoblabe, M. novacekii, M. viridis*, and *M. wesenbergii*) were proposed to be reclassified as a single bacterial species (*M. aeruginosa*) (17). These five *Microcystis* strains showed 16S rRNA gene sequence identities higher than the usual cutoff value used to define bacterial species (>97%), DNA-DNA genome sequences hybridization values were also higher than the cutoff (>70%), and colony morphologies are generally similar (17, 26). Other studies showed that *M. aeruginosa* morphospecies together with other morphospecies are a single species complex with average nucleotide identity (ANI) values higher than 95%, which is consistent with hybridization values higher than 70% – a standard rule in bacterial species delineation (10, 14, 30). Despite the high similarity in their core genomes, *Microcystis* are diverse in their gene content, resulting in large accessory genomes that can harbour genes related to the biosynthesis of harmful toxins or secondary metabolites (10, 14, 31). *Microcystis* classification based on accessory genes (*e.g*. toxins and polysaccharides) has also been proposed (14, 28).

In this study, we present a population genomics analysis using 33 newly sequenced genomes (of which 19 were resequenced after several years in laboratory culture) belonging to seven *Microcystis* morphospecies isolated mainly from Brazil and Canada over a 15-year period. We aimed to investigate the coherence of *Microcystis* morphospecies using phylogenomic and homologous recombination analyses. We identified four *Microcystis* clades corresponding to morphospecies, which are mainly restricted to particular geographical regions, with recombination rates higher within than between them, consistent with a BSC-like model. Meanwhile, *M. aeruginosa* morphospecies are paraphyletic and geographically unstructured, meaning that *M. aeruginosa* may in fact include multiples sub-species. In contrast with the general preference for recombination within clades, we also observed occasional HGT between clades. Many of these cross-species HGT events may be involved in local adaptation. Finally, we studied the profiles of genes related to the biosynthesis of secondary metabolites (such as microcystin) to determine if different *Microcystis* clades have a characteristic profile of biosynthetic genes.

## Results

### Phylogenetic coherence of named morphospecies

In order to assess the coherence of named *Microcystis* morphospecies, we sequenced 33 isolates of *Microcystis* (see Table S1 in the supplemental material). These genomes were initially classified into seven different morphospecies (*M. aeruginosa*, M. *flos-aquae*, *M. panniformis*, *M. wesenbergi, M. viridis, M*. sp. and *M. novacekii)*. The size of the *Microcystis* assembled genomes ranged between 3.3 and 4.6 Mb with average GC content of 42.7%. The shared core genome consisted of 1,260 genes and the pangenome contained 16,928 genes. We supplemented these newly sequenced genomes with 26 additional *Microcystis* genomes downloaded from GenBank and compared them using three measures of genetic similarity: phylogeny, hierBAPS clustering and ANI (Average Nucleotide identity). Most of our genomes have pairwise ANI values > 95% with some exceptions: 199 out of 6241 pairwise comparisons have values between 93 and 94%, mostly involving comparison with Ma_AC_P_00000000_S299 (see Fig. S1, and Table S2 in the supplemental material). ANI values >95% are generally considered to include members of a single species. However, the *Microcystis* genomes do not constitute a single homogenous ANI cluster; rather, significant substructure is evident (Fig. S1).

To explore this substructure, we built a core genome phylogeny using 152 conserved genes also present in an outgroup (Fig. 1) and also clustered the aligned core genomes using hierBAPS (32). Of the 33 newly sequenced genomes, 19 were resequenced after several years in culture. The resequenced genomes differed from their ancestor by an average of 48 point mutations, and always clustered with their ancestor in the phylogenetic tree, suggesting that evolution in the laboratory had little impact on the structure of the phylogeny (Table S3). Additionally, we isolated and sequenced a single colony from one of the batch cultures. This colony genome (S217Col) clustered on the phylogeny with its parent culture (S217Cul) with a phylogenetic distance of zero (Fig. 1), suggesting that a single colony is representative of the entire culture.

**Figure 1.**
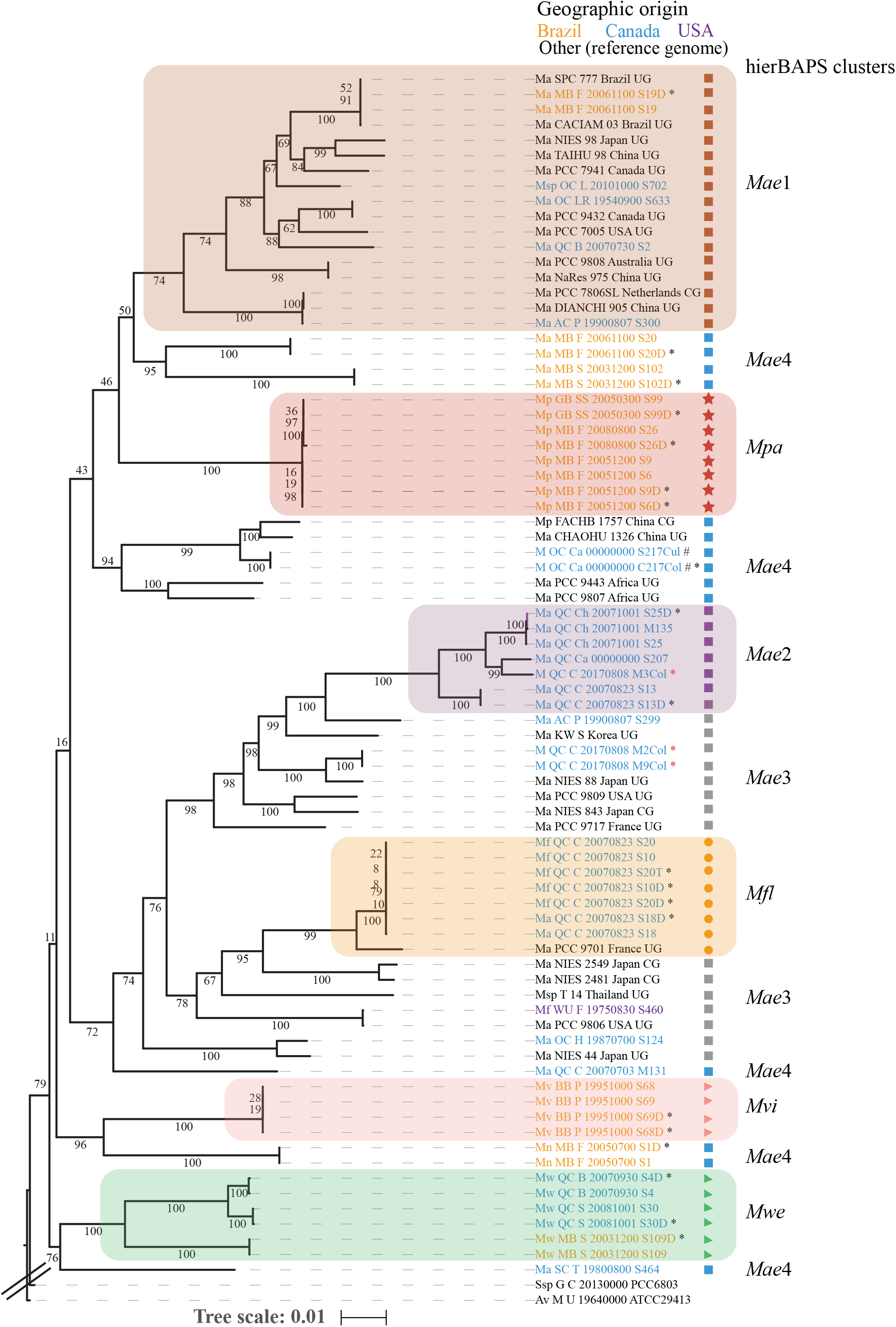
Phylogenetic tree of 53 Brazilian, Canadian and USA *Microcystis* genomes and 26 globally sampled reference genomes. A core genome of 152 homologous genes shared by 79 *Microcystis* genomes and the outgroups (*Anabaena variabilis* ATCC 29413 and *Synechocystis* sp. PCC6803) was identified using software Roary (77). The eight hierBAPS clusters are highlighted in colored boxes. The genomes from the same isolate at a different time have letter “D or T” at the end of their names and are indicated with a black asterisk. The genomes from a bulk culture and a single colony from the same cultured are indicated with a hash. The three genomes from uncultured colonies from Lake Champlain (Quebec, Canada) are indicated with a red asterisk and “Col” at the end of their name. The font colors indicate the geographical origin (Brazil: orange, Canada: blue, USA: purple, other: black). The abbreviated hierBAPS cluster names *Mpn, Mwe, Mfl, Mvi* and *Mae* correspond to *M. panniformis*, *M. wesenbergii M. flos-aquae, M. viridis* and *M. aeruginosa*, respectively. The tree bar scale indicates number of nucleotide substitutions per site.

The hierBAPS analysis yielded eight groups (*Mpa, Mfl, Mvi, Mwe, Mae1, Mae2, Mae3* and *Mae4*) which are indicated alongside the phylogenetic tree (Fig. 1). In general, hierBAPS clusters were consistent with the phylogeny, with some exceptions likely due to the susceptibility of hierBAPS to long-branch attraction (32, 33). Four out of eight hierBAPS clusters corresponded to monophyletic groups in the phylogeny and named morphospecies: *M. panniformis (Mpa), M. wesenbergii (Mwe), M. viridis (Mvi*) and *M. flos-aquae (Mfl)*, while the other clusters contained multiple named morphospecies (Fig. 1). The *M. aeruginosa* morphospecies was paraphyletic and distributed across four hierBAPS clusters (Mae1, *Mae2, Mae3* and *Mae4*), two of which (*Mae1* and *Mae2*) were monophyletic clades. Based on these phylogenetic and population structure analyses, *Microcystis* appears to comprise six well-defined monophyletic groups (*Mpa, Mfl, Mvi, Mwe, Mae1, Mae2*), four of which are congruent with the morphospecies (*Mpa, Mfl, Mvi* and *Mwe)*. We noted that these last four monophyletic morphospecies tended to be fairly specific for a particular geographic location (either Canada or Brazil), suggesting possible local adaptation, local and recent clonal expansion, or reduced migration of these lineages. Even if *Microcystis* are generally closely related at >95% ANI, there is clear and significant substructure within the genus.

We also investigated the pangenome content within each of eight hierBAPS clusters. The four clusters corresponding to *M. panniformis, M. wesenbergii, M. viridis* and *M. flos-aquae* had highly conserved core genomes (between 75% and 96% of genes shared by all members of the cluster) while *M. aeruginosa* morphospecies had much smaller core genomes (between 53% and 64% of genes shared by all members; see Table S4 in the supplemental material). This is consistent with the four monophyletic morphospecies representing coherent clusters of genomic similarity, with the paraphyletic *M. aeruginosa* being an amalgam of high genetic diversity and variable gene content.

### Higher homologous recombination rates within than between clusters supports a BSC-like concept

We next asked if homologous recombination could explain the cohesion of the monophyletic groups. To address this question, we estimated the ratio of homologous recombination to mutation rates (*r/m*) within and between eight hierBAPS groups. We found that the four monophyletic morphospecies (*M. panniformis, M. wesenbergii, M. novacekii* and *M. flos-aquae*) all have *r/m* ratios 2-3X higher within than between clades (Fig. 2). Recombination rates were generally low for *M. aeruginosa* both within and between clades (Fig. 2). When the resequenced genomes (from a second time point of the same culture) were excluded, the *r/m* tended to increase within clades (see Table S5 in the supplemental material), presumably because mutation but not recombination took place in the culture. Overall, these results suggest that the cohesion of four monophyletic groups could be driven or reinforced by preferential recombination within versus between groups, consistent with a BSC-like model of speciation. Conversely, the other four groups, consisting of *M. aeruginosa*, appeared to engage in relatively little recombination and thus defied delineation based on the BSC.

**Figure 2.**
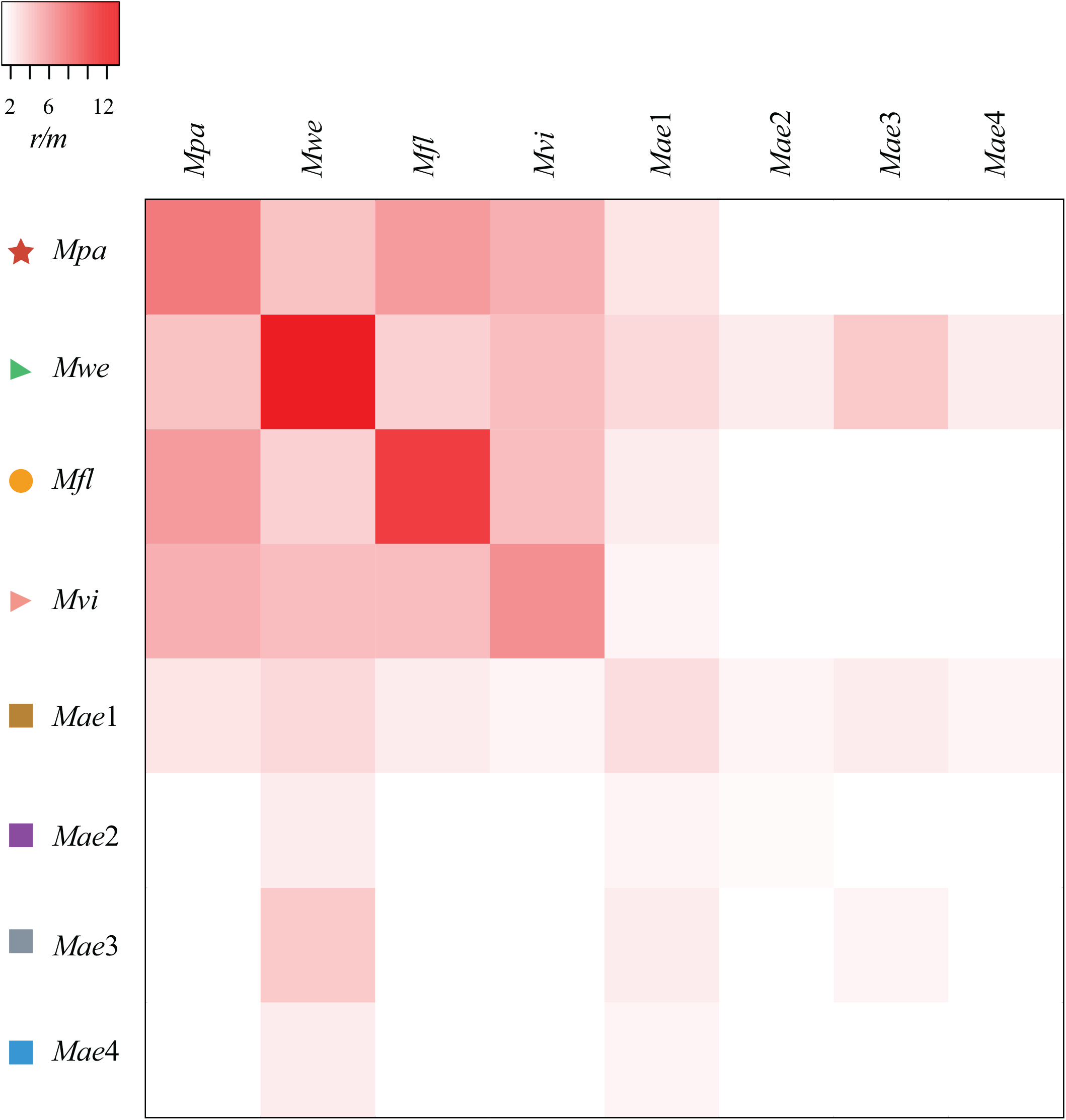
Relative contribution of recombination/mutation (*r/m*) within (diagonal) and among eight *Microcystis* hierBAPS clades. The *r/m* estimation included the replicates genomes (from a second time point of the same culture). The hierBAPS clusters are represented with the same symbols and abbreviations used in Fig. 1. See Table S5 for the *r/m* values with and without replicate genomes.

### Frequent local horizontal gene transfer (HGT)

From the core genome phylogenetic tree, certain monophyletic clades showed strong geographic preferences. For example, *M. flos-aquae* was found almost uniquely in Canada, while *M. panniformis* was found only in Brazil (Fig. 1). In contrast, *M. wesenbergii* is found in both Canada and Brazil (Fig. 1), suggesting that morphospecies can be stable in ways that transcend geography boundaries. As previously observed in *Vibrio*, different species in the same geographic region may exchange genes, possibly leading to local adaptation (7). To identify potential locally adapted *Microcystis* genes, we screened gene trees for instances where two different named morphospecies (which formed distinct monophyletic groups in the species tree; Fig. 1) clustered together in the same monophyletic group (with bootstrap support >90%), consistent with cross-species HGT. As we were particularly interested in local HGT, we identified monophyletic groups of two distinct species, all isolated from the same region (*i.e*. Canada or Brazil, but not both). We screened a total of 25,157 core and accessory genes from 79 *Microcystis* genomes (53 reported here, including colonies and resequenced isolates, and 26 previously published). We considered 12,084 informative gene trees (that included 4 or more leafs). Of these trees, 590 (4.9% of the total) showed a pattern of non-local HGT (with Canadian and Brazilian isolates grouping together in the same well-supported clade), whereas slightly more (923 genes; 7.6% of the total) were consistent with local HGT. This suggests that geography, and possibly local adaptation, is an important factor in shaping rates of HGT. Local HGT events, on average, appear to be more recent than non-local events: in 80 out of 923 local HGTs, the phylogenetic distances within the recombined clade were equal to zero (Table 1), suggesting relatively recent HGT in local compared to non-local events (Fisher’s exact test, Odds ratio = 1.90, *P* = 0.004). Local HGTs are also much more likely to be functionally annotated, compared to non-local HGTs which involve mostly hypothetical genes (Table 1; Fisher’s exact test, Odds ratio = 2.99, *P* < 2.2e-16). While these differences could have many possible explanations, we speculate that non-local HGT events are enriched in phages and other poorly annotated mobile or selfish genetic elements, while local HGTs involve metabolically or ecologically relevant genes, which are more likely to have been studied and annotated. Consistent with this explanation, the non-local pangenome is dominated by genes involved in DNA replication, recombination and repair (COG category L, Fig. S2; *X*^2^ test, *P* < 0.05 after Bonferroni correction for multiple hypothesis testing), which is suggestive of self-replicating and recombining mobile elements (Fig. S2 and Data Set S1). Overall, these results suggest that local HGT events are relatively recent (and thus more frequently observed) and possibly more ecologically relevant (and less “selfish”) than non-local HGTs.

**Table 1.** Breakdown of horizontally transferred genes by geography.

|  | Local HGT | Non-local HGT |
| --- | --- | --- |
|  | (Canada-only or Brazil-only clades) | (Canada-Brazil clades) |
| Hypothetical | 474 | 448 |
| Known Function | 449* | 142 |
| Phylogenetical distances equal to zero | 80* | 28 |
| Total | 923 | 590 |

\* denotes a significant enrichment among local vs. non-local HGT events (Fisher's exact test, $P <$ 0.01).

Figure 3 illustrates a few noteworthy examples of local HGT events. For example, the phylogenetic trees of two neighboring genes encoding the *hicA-hicB* toxin-antitoxin system showed phylogenetic distances almost equal to zero (between 0 and 0.0008 substitutions per site), clustering Brazilian genomes of three different hierBAPS clusters into a single group (Fig. 3A and B), whereas these clusters are well-separated on the species tree (Fig. 3E). This suggests that the toxin-antitoxin system has been subjected to recent cross-species HGT in Brazil. The *hicAB* module is a mobile element that has been previously described in bacteria, archaea, plasmids and phages (34, 35) and at least 31 *hicB* antitoxins and 21 *hicA* toxins have been reported in *M. aeruginosa* (36). The *hicAB* module seems to act as a phage defense system, arresting cell growth in response to phage infection (36).

**Figure 3.**
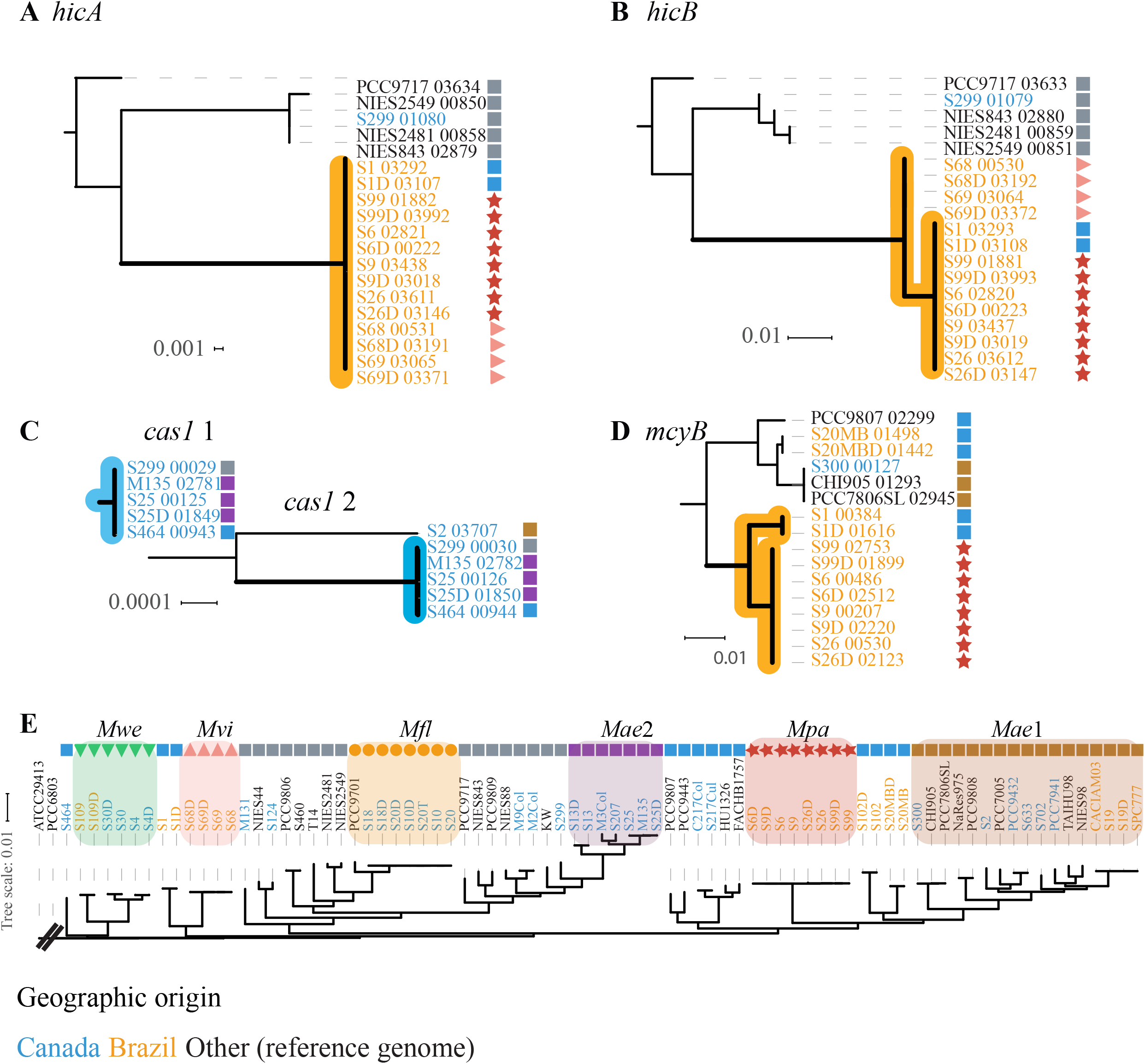
Phylogenetic trees of accessory genes with an inferred history of local HGT. a) *hicA*, b) *hicB*, c) two Cas1 genes, d) *mcyB*, and e) the core genome phylogeny. The symbols represent their corresponding hierBAPS clades. The font colors of the names indicate the geographical origin. The clades showing geographical signatures (local HGT) are highlighted in orange (Brazil) and blue (Canada). The bars below the trees indicate units of nucleotide substitutions per site.

Another two genes encoding CRISPR *cas1* endonucleases also showed a signature of recent local HGT, with phylogenetic distances almost equal to zero (between 0 and 0.0001), clustering Canadian genomes in a single clade (Fig. 3C). These two genes are neighbors located on the same contig, flanked by a hypothetical gene and a CRISPR-associated endoribonuclease (cas2). We also identified local HGT events involving other toxin-antitoxin genes, cyanotoxins (such as *mcyB;* Fig. 3D), endonucleases, and others (Data Set S1).

### Clade-specific profiles of biosynthetic gene clusters

Having shown examples of cyanotoxin genes being involved in local HGT (Fig. 3D; Data Set S1), we sought to more broadly characterize the distribution of cyanotoxins and other biosynthetic gene clusters across *Microcystis* clades. Specifically, we asked whether *Microcysti*s clades or morphospecies tended to have a characteristic profile of biosynthetic genes, despite potentially rapid gain and loss of these genes. The biosynthesis genes of secondary metabolites are usually found in gene clusters (37, 38). *Microcystis* species can synthesize a variable number of secondary metabolites, many of which are toxic to humans and other animals (39, 40).

We identified 34 known secondary metabolite gene clusters within all the *Microcystis* genomes using the software AntiSMASH (Fig. 4; Table S6). AntiSMASH identifies these genes based on a protein database and NRPS (Nonribosomal Peptide Synthetases) and PKS (Polyketide synthase) domain analysis (41). Eight out of 34 secondary metabolic gene clusters were present and complete in at least one *Microcystis* genome (Fig. 4).

**Figure 4.**
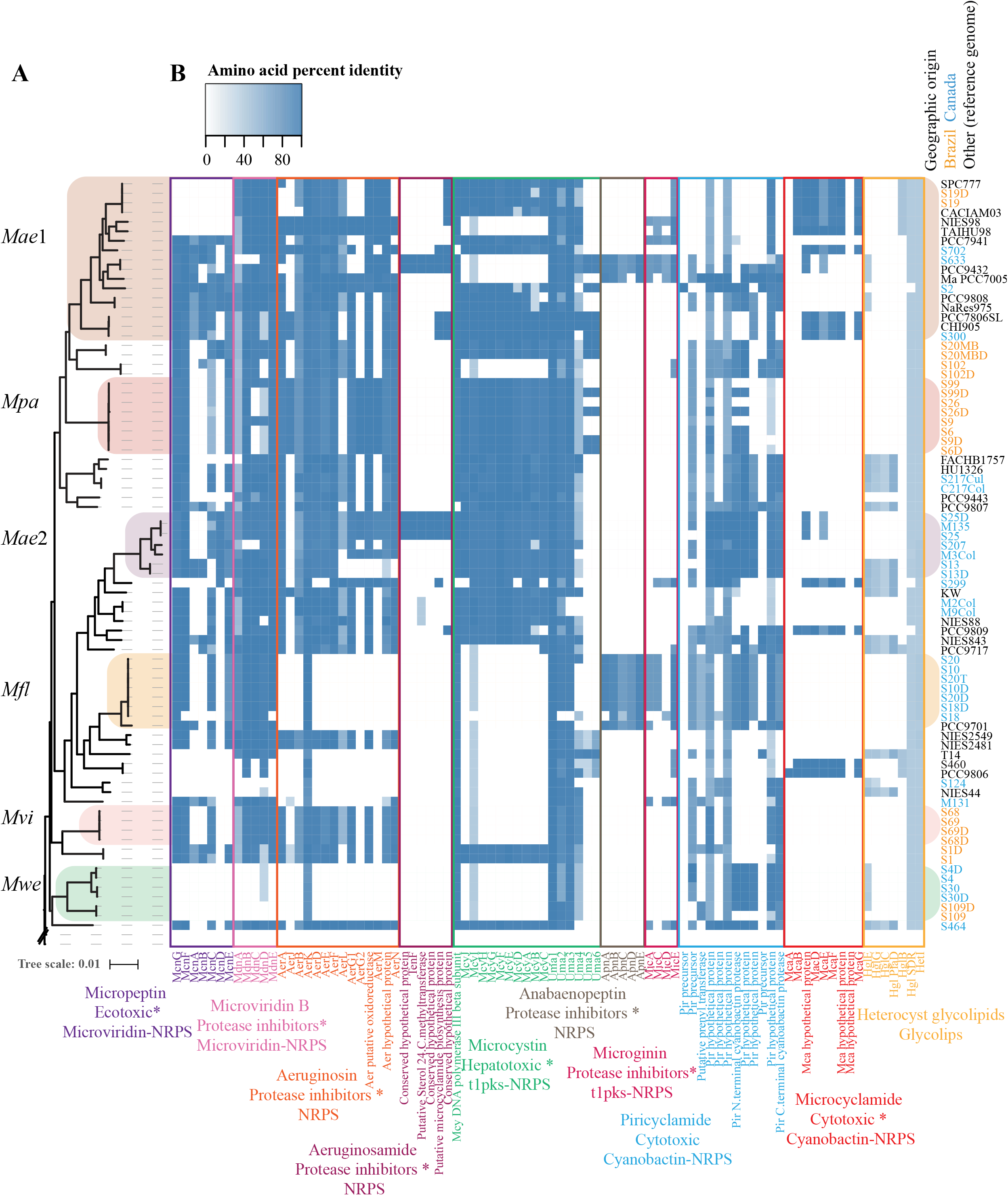
Distribution of biosynthetic gene clusters across *Microcystis*. **a.** Phylogenetic tree of 53 Brazilian, Canadian and USA *Microcystis* genomes and 26 reference genomes. **b.** Presence and absence of the genes encoding secondary metabolites in each *Microcystis* genome shown as a heatmap. Rows and columns represent the genomes and genes, respectively. The presence and absence of genes are indicated in blue and white, respectively. The shade of blue increases with the amino acid similarity to the reference database. The ten biosynthetic clusters are enclosed by colored rectangles and their names appear at the bottom of the figure.

We observed that *Microcystis* genomes lacking the microcystin cluster (*mcy*) usually contained another gene cluster instead. For example, *M. flos-aquae* lacked *mcy* but instead had genes related to the biosynthesis of anabaenopeptins (*apn*) (Fig. 4). However, other studies have found microcystin-producing strains of *M. flos-aquae* (42), suggesting that the genomes reported here likely undersample the diversity of biosynthetic gene clusters present in nature. Consistent with relatively high diversity within clades, genomes within the same clade tended to have similar, but non-identical profiles of gene clusters, with *M. aeruginosa* clades being among the most diverse. *M. aeruginosa* clades also tended to have a high coding potential for toxins, including the complete *mcy* and *mdn* (microviridin B) gene clusters. However, some genomes in *M. aeruginosa* sub-clade *Mae1* lacked the *mcy* genes, consistent with loss or HGT. In contrast, *M. wesenbergii* encoded relatively few biosynthetic gene clusters, consistent with previously reported low toxin production and microcystin gene absence (42–45).

Viewed in aggregate, these biosynthetic gene clusters are part of the *Microcystis* accessory genome. However, certain gene clusters are core to specific monophyletic groups. In *M. flos-aquae* for example, *anp* genes were always present (core) and *mcy* genes were absent. These group-specific core gene clusters could provide potential niche adaptations and ecological distinctness. On the other hand, certain biosynthetic genes such as *mcyB* (Fig. 3D) are exchanged across species boundaries. Thus, biosynthetic genes may contribute to both species-specific and species-transcending adaptations (46).

## Discussion

In this study, we investigated the correspondence among morphospecies and genome-informed species definitions using dozens of *Microcystis* isolates from both Northern and Southern hemispheres, primarily from Canada and Brazil. We assessed the genomic cohesion of *Microcystis* clades within *Microcystis* species by measuring the genome similarities (phylogeny, hierBAPS clustering and ANI values) and homologous recombination within and between clades.

We found that *Microcystis* genomes used in this study together with the reference genomes fell into a single genomic complex (ANI values higher than 95%). Previous studies have suggested a universal cutoff of 95% ANI as adequate for species delineation. These studies described a genetic discontinuity or bimodal distribution with peaks higher than 95% (intra-species) and lower than 83% (inter-species), but the mechanism for this discontinuity is unclear, and it is difficult to exclude sampling bias as a reason for the discontinuity (47, 48). We observed that within a 95% ANI cluster of *Microcystis*, there is substantial genetic substructure, potentially containing distinct species or sub-species. Four of the eight sub-clusters we identified corresponded to named morphospecies, while the others were mostly composed of genomes classified as *M. aeruginosa* morphospecies. We concluded that *M. aeruginosa* is paraphyletic with a mixed geographical pattern, while the morphospecies *M. flos-aquae, M. panniformis, M. wesenbergii, M. viridis* consisted of well-defined clades within *Microcystis* species complex (Fig. 1).

What are the mechanisms that can explain the genetic structure within *Microcystis*? Ecological selection (the ESC), barriers to gene flow (BSC-like), and biogeography (allopatric divergence) could all play a role. Previous studies based on a smaller sample of *Microcystis* genomes (10), or marker genes (14, 15) suggested that there are few if any biogeographic barriers in *Microcystis* to migrate, leading to a globally mixed population (49, 50) – and that *Microcystis* should be defined as a single species (17, 18). Consistent with this, *M. aeruginosa* is globally distributed and paraphyletic in our core phylogeny. However, the four monophyletic morphospecies tend to be geographically restricted, possibly due in this case to limited migration (at least on recent time scales) and/or local adaptation. The four morphospecies could also represent short-lived clonal expansions, or biases due to incomplete sampling. However, to minimize bias, our sampling was performed repeatedly over 15 years, with similar methods in both Brazil and Canada. We also inferred more frequent homologous recombination within than between morphospecies (Fig. 2), consistent with a BSC-like model maintaining genetic distinctness (51–53). While morphospecies coherence may thus be maintained by barriers to gene flow, we suspect that morphospecies divergence was initiated by selection for ecological distinctness (1). Although the precise ecological differences between morphospecies are unknown (45, 54), we found that each of the four monophyletic morphospecies had a distinct core genome and distinct profile of biosynthetic gene clusters (Fig. 4) – both of which could provide potential ecological adaptations. Further experimental study will be required to fully test the hypothesis of ecological distinctness among morphospecies.

The BSC-like model requires more frequent recombination within than between species, but also allows occasional recombination of “globally adaptive” genes across species boundaries. Similar to previous observations in *Vibrio* (7), we inferred a significant proportion of cross-species HGT events occurred within the same geographic location, suggesting local environmental adaptation. Local HGT events tend to be phylogenetically more recent than non-local events, suggesting that they occur at relatively higher frequency. Local HGTs are also enriched in genes of annotated (non-hypothetical) function, including cyanotoxin genes such as microcystin (mcyB) and cyanopeptolin (*mcnC, mcnF* and *mcnG*). Microcystin genes are likely of ancient origin in cyanobacteria (55) and it has been suggested that they subsequently experienced significant homologous recombination and positive selection (56, 57). Microcystin genes also show biogeographic patterns. For example, the *mcyD* gene has distinctive alleles found in Japanese *Microcystis* isolates but not elsewhere (58). Our inference of local, cross-species HGT of toxin genes further supports the idea that they may be locally adapted.

Genes involved in phage defense systems were also involved in local HGT events, suggesting that local adaptation could be driven by local viruses. First, the *hicAB* operon appears to have been shared among at least three distinct hierBAPS clusters (*Mvi, Mpa*, and *Mae4*) in Brazil (Fig. 3), and has previously been suggested to be involved in phage defense and prone to HGT (34, 36). Second, *cas1*, which encodes the most conserved protein in the CRISPR–Cas defense system (59), appears to have been exchanged among three distinct clusters of *M. aeruginosa* in Canada (Fig. 3). This is consistent with previous evidence suggesting that CRISPR–Cas genes are subject to HGT and natural selection (59, 60). Thus, local HGT could promote adaptation to local phages.

Taken together, our results resolve some of the longstanding confusion surrounding *Microcystis* species and suggest new avenues for future research. While all *Microcystis* genomes sampled to date are monophyletic and closely related, there is significant genetic substructure suggesting the existence of several distinct species. The distinctiveness of these species appears to be maintained by barriers to gene flow, consistent with a BSC-like model (1, 5). Whether gene flow barriers are mainly geographic, genetic, or ecological is a subject for future investigation. While different *Microcystis* species appear to inhabit different niches, as evidence by geographic preferences and distinct profile of biosynthetic gene clusters, the nature of their ecological distinctiveness should also be a subject of future field and laboratory studies.

## Materials and methods

### Genome sequencing, assembly and binning

Over the past 15 years, we collected 30 *Microcystis* isolates from Brazil, Canada and the United States. The *Microcystis* strains were initially characterized as morphospecies based on their colony morphology, according to Komárek (27, 61, 62). Seven morphospecies were identified: *M. aeruginosa*, M. *flos-aquae, M. panniformis, M. wesenbergi, M. viridis, M*. sp., and *M. novacekii*. We performed the DNA extraction for these strains between 2006 and 2017 (see Table S1 in the supplemental material). The 30 *Microcystis* genomes were sequenced using the Illumina HiSeq 2500 platform with 125bp paired-end reads. The genomic Illumina libraries (with average fragment size 360bp) were prepared using the NEB (New England Biolabs®) low input protocol. We also sequenced the DNA of four single *Microcystis* colonies isolated manually under the microscope. Three of these colonies were new isolates recovered from Lake Champlain (Quebec, Canada) in 2017 without culture, while the fourth came from a culture that was also sequenced (in bulk) in this study, for a total of 33 new sequenced *Microcystis* genomes. Before DNA extraction and sequencing, each colony was washed 10-15 times with Z8 medium using a micropipette. DNA extraction was performed directly on each colony using the ChargeSwitch® gDNA Mini Bacteria Kit.

Of the 30 *Microcystis* isolates, 19 had been maintained in culture for several years until 2017. Thus, we extracted DNA and re-sequenced from these 19 cultures in 2011, 2016 and 2017 to check for contamination and mutations in the *Microcystis* genome over time and differences between culture and colony sequences (Table S1). Together, the *Microcystis* genomes, of which 14 were sequenced once, 18 twice, and one three times comprised a total of 53 genome sequences.

Sequences from cultures and colonies were assembled with the software IDBA-UD v1.1.3 (63), producing contigs belonging to both *Microcystis* and associated heterotrophic bacteria, which are naturally associated with *Microcystis* (64, 65). The software Anvi’o v3.0 was used to filter, cluster and select the contigs belonging to *Microcystis* (66). The associated bacterial genomes will be described in a forthcoming manuscript. To cluster contigs into genomes, we used tetra-nucleotide frequency, GC content, and taxonomic annotation based on the Centrifuge software, as implemented in Anvi’o (66, 67). The gene prediction and annotation were done using Prodigal v2.6.3 and Prokka v1.12 packages, respectively (68–70). The raw reads and the *Microcystis* genomes contigs are available in GenBank under Bioproject number PRJNA507251.

### Phylogenomic analysis

A core genome of 152 single copy genes shared by 79 *Microcystis* genomes (53 and 26 genomes reported here and previously, respectively) and two outgroups (*Anabaena variabilis* ATCC 29413 and *Synechocystis* sp. PCC6803) was identified using the software Roary and blastn-all. First, a core genome for the 79 *Microcystis* genomes (minimum value of 90% amino acid identity) was identified using Roary. The outgroups were initially excluded due to their high divergence from *Microcystis*. To identify the homologous genes in the outgroups, blast-all was used (Blastp similarities higher than 60%). The common core genes between Roary and blast-all were selected and used to create a core gene alignment. Each homologous gene was aligned separately using muscle (71). The concatenated and degapped alignment of length 129,835 bp was used for building a phylogenetic tree in RAxML v8.2.4, using the GTRGAMMA model, with 100 bootstraps (72). Using the same method, another core phylogenetic tree was inferred without the previously published reference genomes. This concatenated core-alignment comprised 222 genes (211,589 bp degapped) (see Fig. S3 in the supplemental material).

### Clustering analysis of *Microcystis* genomes

A multiple genome alignment of the 79 genomes (53 and 26 genomes reported here and the references, respectively) was performed using Mugsy 1.2.3 (73) (see Table S1 in the supplemental material). The DNA core-alignment was extracted from the Mugsy output using a python script (625,795 bp long). The core-alignment was used to perform genetic population structure and cluster analysis with hierBAPS (The hierarchical Bayesian Analysis of Population Structure) (32). We used as input parameters two clustering levels and an expected number of cluster (k) equals to 10, 20 and 40. HierBAPS delineates the population using nested clustering. In this method, rare genotypes (distantly related to better sampled clades) often cluster together due to long-branch attraction (32). As a result, HierBAPS clusters could be incongruent with the phylogeny inferred by maximum likelihood, which is less sensitive to long-branch attraction.

### Pairwise nucleotide identity between *Microcystis* genomes

The 33 newly sequenced genomes and 26 previously published were compared using a python module (average_nucleotide_identity.py) to estimate the Average Nucleotide Identity by Mummer and by Blast (ANIm and ANIb values between genome pairs) (see Fig. S1 in the supplemental material) (https://github.com/widdowquinn/pyani) (74). Bacterial genomes with DNA-DNA hybridization (DDH) of at least 70%, are considered as the same species and usually show values of ANI >95%. Hence, a cutoff higher than 95 for the ANI values between genome pairs is used to identify genomes within the same genomic cluster or species (30, 47, 75). The pairwise identities were plotted using the R package ggplot (http://ggplot2.org/) and the function heatmap2 (76).

### Pangenome analysis in *Microcystis* genomic clusters

Pangenomes were inferred for each *Microcystis* genomic cluster with more than 2 genomes (see Table S2 in the supplemental material). A global pangenome estimation was generated using all the *Microcystis* genomes excluding the 2 shorter genomes Ma_AC_P_00000000_S299 and Ma_QC_C_20070823_S18). Those genomes were excluded of the pangenome analyses because of their reduced size compared to the average size (10% and 30% for Ma_AC_P_00000000_S299 and Ma_QC_C_20070823_S18, respectively). Ma_QC_C_20070823_S18 showed the lowest coverage (28X) and Ma_AC_P_00000000_S299 appeared to be contaminated with another cyanobacterium (*Anabaena*).

The software Roary v3.12.0 was used to generate the pangenomes. Prior to the execution of Roary the genomes were automatically annotated using Prokka v1.12 (70). The genomes in GGF3 format generated with Prokka were used as input to Roary. Roary was executed using a minimum percentage of amino acid identity of 90% for blastp, which was set up according to the similarities in the genomes higher than 94% and Roary recommendations. MultiFasta alignment of the core genes were created using PRANK v150803 implemented in Roary (77).

We also did a pangenome analysis to find homologous and accessory genes within 53 *Microcystis* genomes from Brazil, Canada, USA (including the shorter ones), and the 26 reference genomes from the NCBI database. Roary allowed us to identify 370 clusters of homologues (shared in 79 genomes) and 1059 (shared in 76 up to 78 genomes). Roary also identified 23,728 accessory genes or genes shared by less than 76 genomes.

### SNPs and deletion identification between duplicates genomes

The calling of SNP variants (Single Nucleotide Polymorphism) was done using snippy v3.2 (https://github.com/tseemann/snippy) with default parameters. SNPs between duplicate genomes were identified using one of genomes as reference.

### Homologous recombination rates across *Microcystis* genomic clusters

Using the 53 newly sequenced *Microcystis* genomes, we investigated rates of homologous recombination within and between genomic clusters or across the phylogenetic tree. To do this, we estimated the relative effect of recombination versus mutation (*r/m*) rates using ClonalFrameML v1.11-3 (78). Briefly, the degapped core genome alignment generated by Mugsy v2.2.1 (1,274,628 bp) (73) was split in several subalignments using pyfasta (https://pypi.org/project/pyfasta/). The subalignments corresponded to the genomic clusters defined using phylogenomic and the population structure analyses. In order to estimate *r/m* between clusters, we also created subalignments for pair of clusters. ClonalFrameML was executed using a bootstrap of 100 replicates (emsim=100). The input phylogenies given in ClonalFrameML were generated using RAXML with bootstrap of 100 replicates (72). The transition/transversion ratios also used as an inputs in ClonalFrameML were estimated using PHYML v3.0 under the model of nucleotides substitution HKY85 (79). ClonalFrameML analyses excluded the two smallest *Microcystis* genomes Ma_AC_P_00000000_S299 and Ma_QC_C_20070823_S18. ClonalFrameML analyses within clusters were also conducted after removing the replicate (resequenced) genomes.

### Identification of horizontally transferred and locally adapted genes

To identify genes transferred across species boundaries, we screened gene trees for instances of two distinct species (hierBAPS clusters in the species tree) clustering together in the same monophyletic group, whereas they are normally distantly related in the species tree. As a signature of local adaptation, we additionally screened for such cross-species HGT events that occurred among two different species from the same country.

For local adaptation analysis, we worked with the clusters generated with Roary and using 53 *Microcystis* genomes from Brazil, Canada, USA, and the 26 reference genomes from the NCBI database. Once the gene clusters were identified with Roary, the alignments of the nucleotide sequences in each gene cluster (core genes present in >75 strains (>95%) and accessory genes) were generated using the MAFFT software v7.271 (80). Maximum likelihood phylogenetic trees for each alignment were inferred using FastTreeMP v2.1.8 and the generalized time-reversible model (GTR) for nucleotide substitution (81). The trees were visualized with graphlan v0.9.7 (82).

The phylogenetic trees that showed local (geographic) adaptation signatures were identified using a Perl script (https://figshare.com/articles/Monophy_screening_tree_files_for_the_detection_of_local_adaptations/7661009) to screen phylogenetic trees and identify monophyletic groups with a particular level of bootstrap support (in our case, 90%). The script also allows monophyletic groups including particular combination of isolates (e.g. from different morphospecies or geographic locations) to be identified, with a given minimal branch length (in our case, this parameter was set to 0) and number of isolates (in our case, 4) within the group. A phylogeny was considered as positive for non-local HGT if Canadian and Brazilian isolates were together in the same clade supported by a bootstrap value > 90%, while a phylogeny positive for the local-HGT showed Brazilian or Canadian isolates, but not both in the same well-supported clade. The phylogenies with a signature of HGT were then manually curated to remove those consisting solely of HGT within a single hierBAPS cluster. Genes in the accessory genome and core genome were functionally annotated using the eggNOG database (83, 84). The full HGT gene set is reported in Data Set S1.

### Inferring secondary metabolic pathways in *Microcystis* genomes

We evaluated the metabolite profiles for individual *Microcystis* genomes using the package antiSMASH v4.0.2. The annotated genomes using Prokka were used as input to AntiSMASH (41). Two additional biosynthetic clusters absent in the AntiSMASH database (Aeruginosamide (NCBI accessions numbers CCH92964-CCH92969) and Microginin (NCBI accessions numbers CAQ48259-CAQ48262)) were added manually. Based on the antiSMASH results, we generated a matrix of presence-absence of genes related to the biosynthesis of secondary metabolites. The matrix was visualized using the R package ggplot and the function heatmap2 (76). All-against-all BLASTP analysis was applied to find the best reciprocal hits between proteins in the database and the proteins in the *Microcystis* genomes (85). The proteins with the best reciprocal hit were extracted; to be considered as present in the database, the amino acid identity had to be > 60%, and > 30 % of the length of the sequences had to be aligned, with an e-value of 10^−5^.

## Acknowledgments

We are grateful to David Bird for assistance isolating strains and maintaining cultures. This work was supported by the Genome Québec and Genome Canada-funded ATRAPP Project (Algal blooms, Treatment, Risk Assessment, Prediction and Prevention). **NT** was funded by a project from the European Union’s Horizon 2020 research and innovation program under the Marie Sklodowska-Curie grant agreement No 656647. Cultures collections (Brazilian and some Canadian strains) were partially obtained and maintained thanks to CNPq and FAPEMIG grants to **AG.** We also want to acknowledge the financial support of the National Research Council. We declare that we have no financial conflicts of interest.

---

**Figure S1. Heatmaps for the hierarchical clustering of 79 *Microcystis* genomes based on ANIm (A) and ANIb (B)** (53 genomes sequenced in this work, plus 26 NCBI genomes published before). Rows and columns represent the genomes. The scale bar indicates the pairwise ANI values. The genomes from the same isolate at a different time point have a letter “D or T” at the end of their names and are indicated with black asterisks. The genomes from colonies and culture from the same isolate are indicated with a hash. The genomes names from colony end with the word Col and are indicated with a red asterisk.

**Figure S2. Functional enrichment analysis in core, accessory, local and non-local HGT gene sets.** Functional annotations were from eggNOG. a) Hierarchical clustering analysis based in the relative abundance for each functional category. b) Functional categories that showed statistically significant (*P* < 0.05) differences in one or more comparison between groups (core, accessory genes, local, or non-local HGT gene sets, defined as described in Methods). The chi-square (*X*^2^) test was conducted using the package STAMP (86). Raw p-values are indicated in black, and after Bonferroni correction in blue. Asterisks denote significant (p < 0.05) differences after correction.

**Figure S3. Phylogenetic tree of 53 Brazilian, Canadian and USA *Microcystis* genomes.** A core genome of 222 homologous genes shared by 53 *Microcystis* genomes and the outgroups (*Anabaena variabilis* ATCC 29413 and *Synechocystis sp*. PCC6803) was identified using the software Roary (77). The concatenated core-alignment (222 genes - 211589 bp long) was used for building a phylogenetic tree in the RAxML program, using the GTRGAMMA model, with 100 bootstrap replicates (72). The genomes from the same isolate at a different time have letter “D or T” at the end of their names and are indicated with a black asterisk. The genomes from colony and culture from the same isolate are indicated with a hash. The genomes names from colonies end with the word Col and are indicated with a red asterisk.

## Supplementary table legends

**Table S1. Genome characteristics of 33 *Microcystis* isolates from Brazil, Canada and USA.** The genomes from the same isolate at a different time point are indicated with black asterisks. The genomes from colonies and culture from the same isolate are indicated with hashes. The genomes names from colonies end with “Col” are indicated with red asterisks.

**Table S2. ANIb and ANIm values corresponding to pairwise comparison between genomes.**

**Table S3. SNP variants between 19 duplicates (resequenced) genomes.** The duplicate (resequenced) genomes are indicated with “D” at the end of their name.

**Table S4. Core genome composition for each hierBAPS clusters and all the genomes.** The core genomes were estimated for each individual hierBAPS cluster using the software Roary. The number of genomes per pan-genome and the ANIm values within clusters are indicated.

**Table S5. The relative effect of recombination versus mutation (*r/m*) within and between hierBAPS clusters.** The *r/m* estimations after removing duplicate genomes are indicated within parentheses and with *.

**Table S6.** Biosynthetic pathways.

**Data Set S1. Genes showing evidence of local and non-local HGT.** The list of genes with local and non-local annotation, phylogenetic distances to the node, and functional annotation with Prokka and eggNOG are indicated.

