## Supplementary material for "Coherence of *Microcystis* species revealed through population genomics": Table S5

Table S5. The relative effect of recombination versus mutation (*r/m*) within and between hierBAPS clusters

| hierBAPS Cluster | Genomes Number | Genomes number no duplicates | *M. panniformis* | *M. wesenbergii* | *M. flos-aquae* | *M. viridis* | *M. aeruginosa 1* | *M. aeruginosa 2* | *M. aeruginosa 3* | *M. aeruginosa 4* |
| --- | --- | --- | --- | --- | --- | --- | --- | --- | --- | --- |
| *M. panniformis*  *Mpa* | 8 | 4 | 7.35 *(11.064) | 3.88 | 5.8 *(6.13) | 4.86 | 2.35 | 1.35 | 1.27 | 1.37 |
| *M. wesenbergii* | 6 | 3 | 3.88 | 13.25 *(13.24) | 3.5 | 4.4 | 3 | 2.33 | 3.6 | 2.04 |
| *M. flos-aquae*  *Mfl* | 6 | 3 | 5.8 *(6.13) | 3.5 | 10.28 *(15.78) | 4.3 | 2.2 | 1.33 | 1.4 | 1.21 |
| *M. viridis*  *Mvi* | 4 | 2 | 4.86 | 4.4 | 4.3 | 6.47 | 1.96 | 1.35 | 1.2 | 1.27 |
| *M. aeruginosa 1*  *Mae1* | 6 | 5 | 2.35 | 3 | 2.2 | 1.96 | 2.92 *(2.92) | 1.73 | 2.16 | 1.75 |
| *M. aeruginosa 2*  *Mae2* | 7 | 5 | 1.35 | 2.33 | 1.33 | 1.35 | 1.73 | 1.48 *(1.48) | 1.31 | 1.33 |
| *M. aeruginosa 3*  *Mae3* | 4 | 5 | 1.27 | 3.6 | 1.4 | 1.2 | 2.16 | 1.31 | 1.88 | 1.12 |
| *M. aeruginosa 4*  *Mae4* | 10 | 6 | 1.37 | 2.04 | 1.21 | 1.27 | 1.75 | 1.33 | 1.12 | 1.3 |

**r/m* rates removing duplicates genomes.
