## Supplementary material for "Coherence of *Microcystis* species revealed through population genomics": Table S4

Table S4. Core genome composition for each hierBAPS clusters and all the genomes.

| hierBAPS cluster | Morpho-species | Number of genomes without replicates |  | Number of genomes per pan-genome | ANIm  Within ANI clusters | Core Genome | Accessory Genome | Total Genes |
| --- | --- | --- | --- | --- | --- | --- | --- | --- |
| *Mpa* | *M. panniformis* | 4 |  | 8 | 99.79 | 4077  (92.19%) | 590 | 4667 |
| *Mwe* | *M. wesenbergii* | 3 |  | 6 | 97.20 | 3136  (75.28%) | 2195 | 5331 |
| *Mfl* | *M. flos-aquae* | 3 |  | 6 | 99.90 | 4143  (96.29%) | 256 | 4399 |
| *Mvi* | *M. viridis* | 2 |  | 4 | 99.88 | 4267  (94.34%) | 396 | 4663 |
| *Mae1* | *M. aeruginosa*  *M.* sp. | 5 |  | 6 | 96.81 | 2617  (59.73%) | 4530 | 7147 |
| *Mae2* | M. aeruginosa | 5 |  | 7 | 97.65 | 2224  (53.50%) | 4495 | 6719 |
| *Mae3* | *M. flos-aquae* | 5 |  | 4 | 95.83 | 2594  (63.69%) | 3654 | 6248 |
|  | *M. aeruginosa* |  |  |  |  |  |  |  |
|  | *M.* sp. |  |  |  |  |  |  |  |
|  | *M. aeruginosa* |  |  |  |  |  |  |  |
| *Mae4* | *M. novacekii* | 6 |  | 10 | 95.05 | 2682  (61.12%) | 7566 | 10248 |
|  | *M. aeruginosa* |  |  |  |  |  |  |  |
|  | *M. aeruginosa* |  |  |  |  |  |  |  |
|  | *M.* sp. |  |  |  |  |  |  |  |
|  | *M. aeruginosa* |  |  |  |  |  |  |  |
| Global pangenome |  | 33 |  | 51 | 95 | (strict core of 1260 shared =>99% of the genomes or a soft core  1833 shared =>95% of the genomes) | 15095 | 16928 |

The two smallest *Microcystis* genomes Ma_AC_P_00000000_S299 and Ma_QC_C_20070823_S18 were excluded from the global pangenome analyses.
