## Supplementary figures and images for "Coherence of *Microcystis* species revealed through population genomics"

### Figure S1

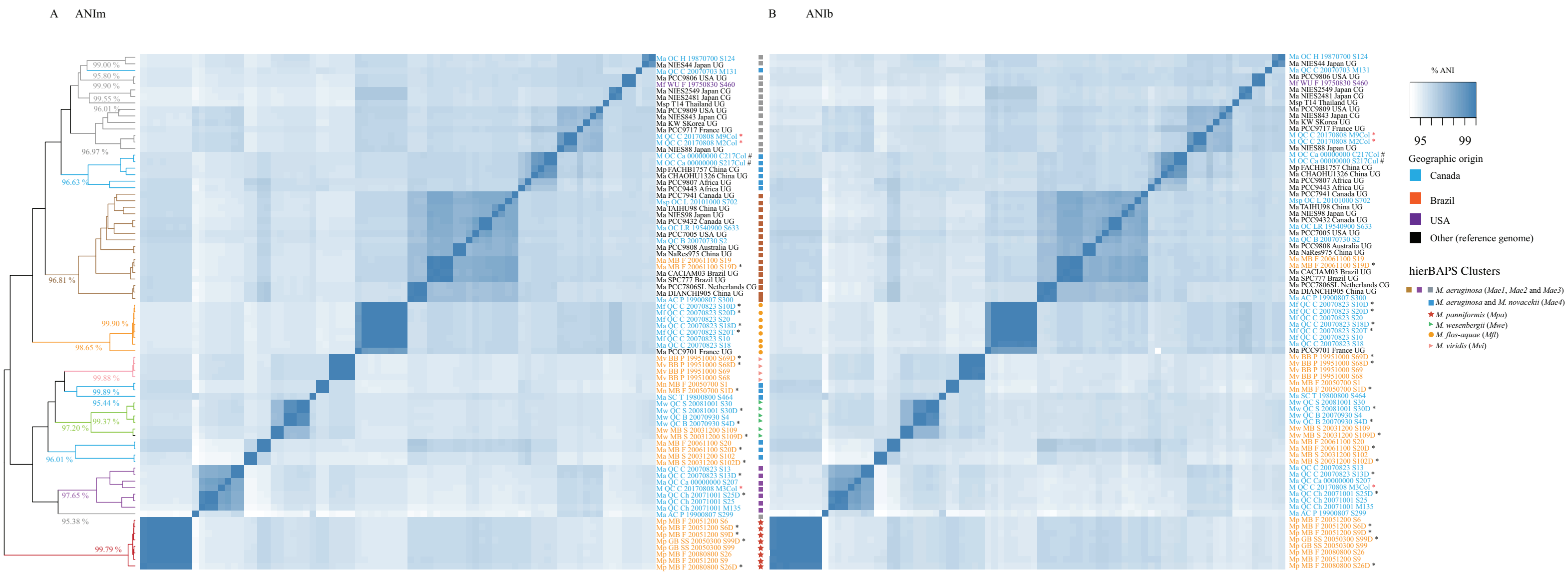

### Figure S2

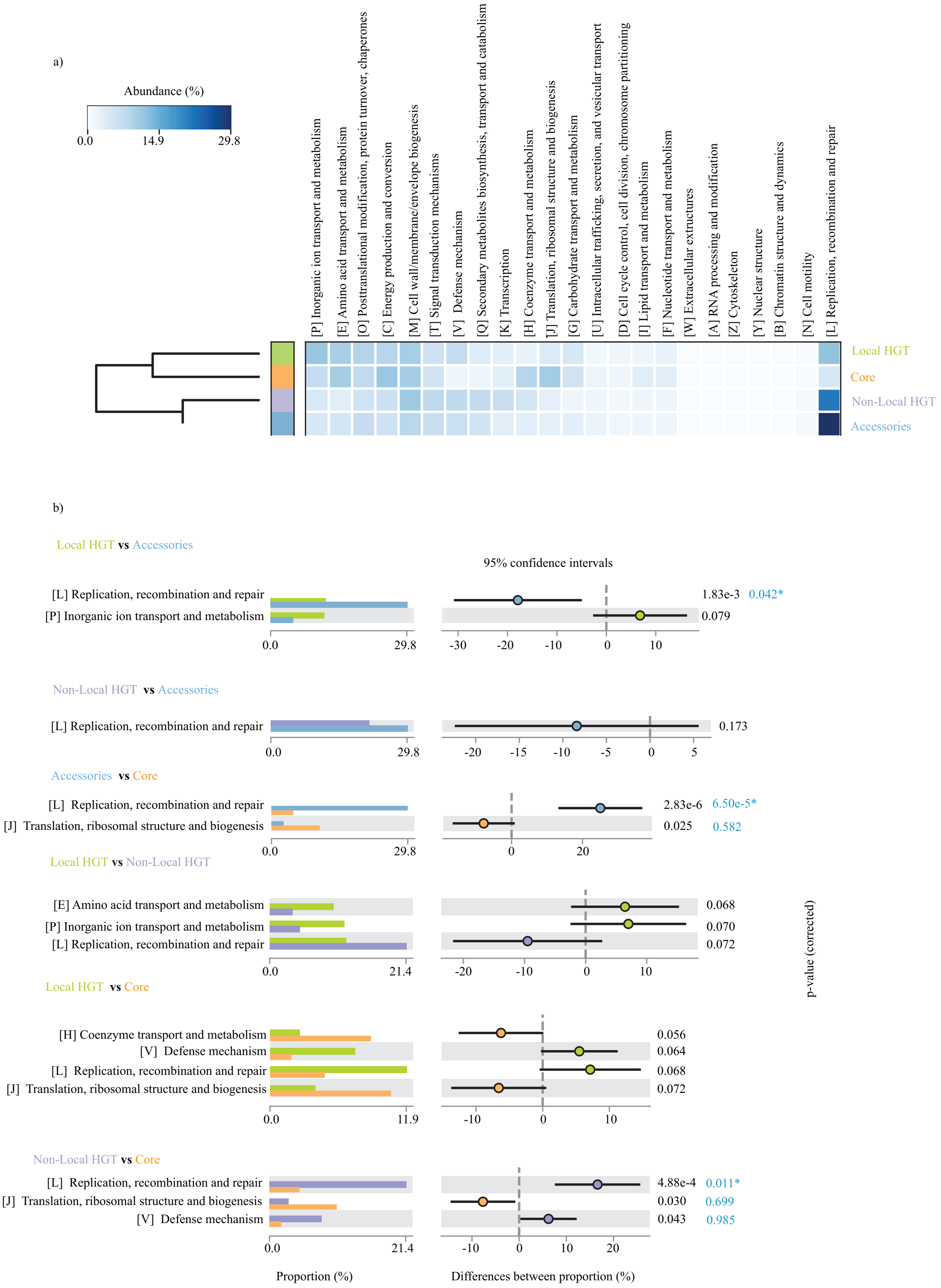

### Figure S3

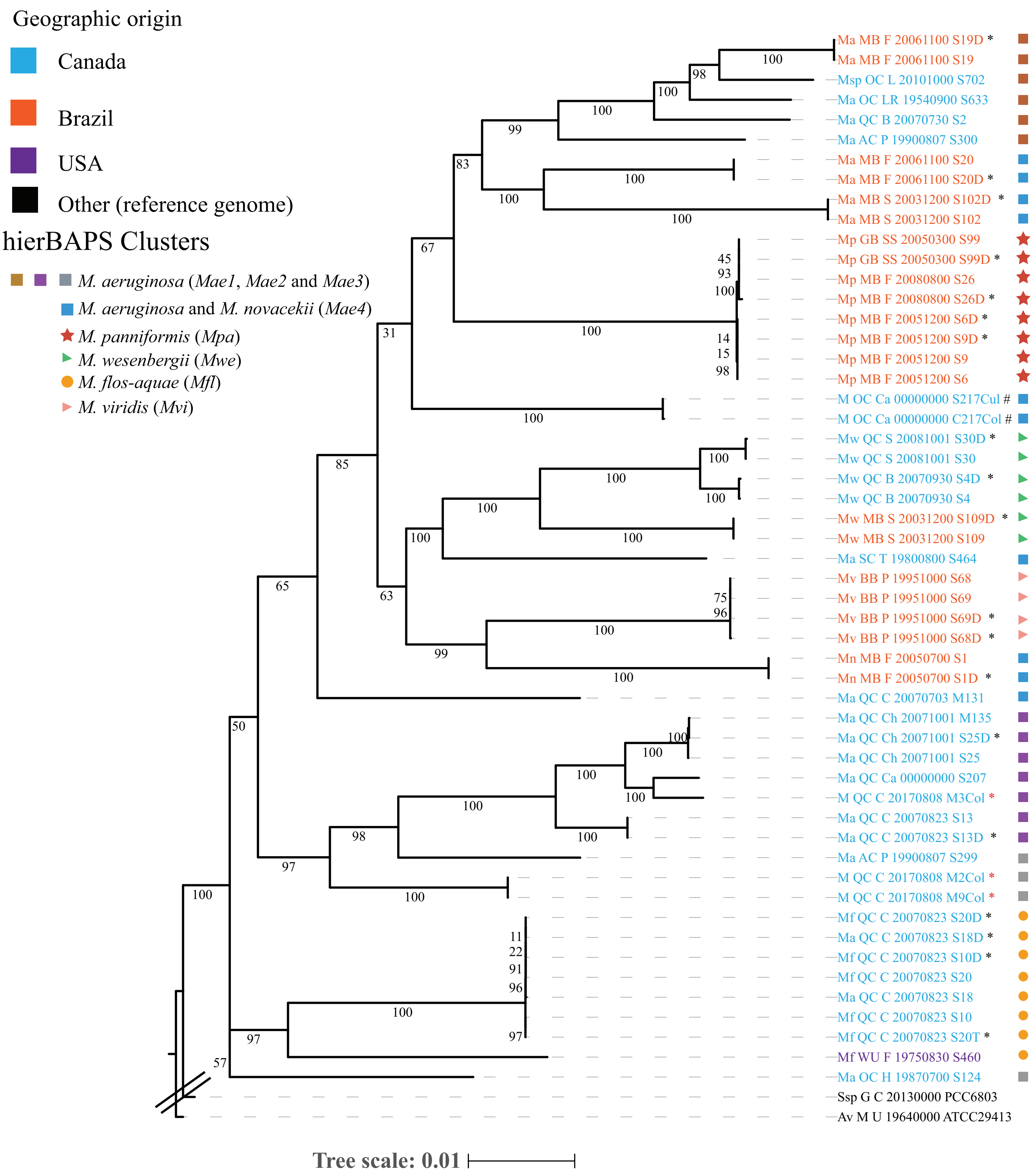
